# Selective and context-dependent social and behavioral effects of Δ^9^-tetrahydrocannabinol in weakly electric fish

**DOI:** 10.1101/269803

**Authors:** Brandon Neeley, Tyler Overholt, Emily Artz, Steven G Kinsey, Gary Marsat

## Abstract

Cannabinoid (CB) receptors are widespread in the nervous system and influence a variety of behaviors. Weakly electric fish has been a useful model system in the study of the neural basis of behavior but we know nothing of the role played by the CB system. Here, we determine the overall behavioral effect of a CB receptor agonist (i.e., Δ^9^-tetrahydrocannabinol, THC) in the weakly electric fish *A. leptorhynchus*. Using various behavioral paradigms involving social stimuli, we show that THC decreases locomotor behavior as in many species and influences the communication and social behavior. Across the different experiments we found that the propensity to emit communication signals (chirps) and to seek social interactions was affected in a context-dependent manner. We explicitly tested this hypothesis by comparing the behavioral effects of THC injection in fish placed in a novel versus familiar social and physical environments. THC-injected fish were less likely to chirp than control in familiar situation but not in novel ones. The tendency to be in close proximity was affected only in novel environments whith control fish clustering more than THC-injected ones. By identifying behaviors affected by CB agonists, our study can guide further comparative and neurophysiological studies of the role of the CB system using weakly electric fish as a model.

## Introduction

Endocannabinoids serve as one of the main retrograde neurotransmitter in the brain [Kreitzer, 2002] and CB receptors are prevalent throughout the nervous system [Svizenska et al., 2008]. They are present across vertebrates [Elphick, 2002; Elphick, 2012], including non-mammalian species (e.g. lampreys [Kettunen et al., 2005], teleosts fish [Cottone et al., 2005; Lam et al., 2006], reptiles [Newman et al., 2007], and birds [Alonso-Ferrero et al., 2006; Stincic, and Hyson, 2008]). The role of the CB system in neural function and its influence on behavior present many similarities across vertebrate species [Dalton et al., 2009; Elphick, 2012]. CB agonist decrease locomotor activity in several vertebrates [Soderstrom et al., 2000; Soderstrom, and Johnson, 2001; Valenti et al., 2005] as it does in mammals [Sañudo-Peña et al., 2000]. Effects on appetite [Soderstrom et al., 2004; Valenti et al., 2005], learning and memory [Soderstrom, and Johnson, 2003] and propensity to produce communication signals [Soderstrom, and Johnson, 2001] are also affected non-mammal as it is in mammals. Of particular relevance to our study, social interactions are enhanced by cannabinoid agonist in several species (e.g. zebrafish [Barba-Escobedo, and Gould, 2012] or mice [Umathe et al., 2009]). The influence of cannabinoids on anxiety is well documented (e.g. [Moreira, and Wotjak, 2010]) and CB agonist can alter the reaction to novel environment (e.g. zebrafish [Connors et al., 2014] or mice [Haller et al., 2004]). In mice, the effect on stress/anxiety and the effects on social interaction seem to be opposite in direction depending on the exact experimental and environmental conditions [Akinshola et al., 1999; Haller et al., 2004; Krug, and Clark, 2015; Navarro et al., 1997]. These studies are among the many studies on mammals detailing the complexity of the behavioral effect of cannabinoid but we are still figuring out which of these effects are generalizable across vertebrates and which are species specific. We can therefore hardly predict which effect will be observed in any given species. In this paper, we aim to detail some of the main effect of CB agonist on the behavior of weakly electric fish; a model system that has led to important advances in our understanding of the neural basis of behavior.

The weakly electric fish has historically been a very useful model system for advancing our understanding of both sensory processing [Allen, and Marsat, 2018; Maler, 2007; Zakon, 2003] and sensory-motor integration [Sawtell, 2017], and can thus be credited with some of the most detailed understanding of how specific behavioral responses can be generated by sensory-to-motor neural pathways [Heiligenberg, 1991; Rose, 2004]. This deep understanding of the neural basis of complex behavior will facilitate linking specific behavioral effect of CBs with the underling neural mechanisms. Furthermore, there are over 200 species of closely related gymnotiformes, each displaying differences in their behavior. They are therefore well suited for comparative studies that could reveal how the CB system has evolved in relation to behavior.

Gymnotiformes vary in social grouping, from gregarious to solitary [Stamper et al., 2010]. *A. leptorhynchus* tends to be found in groups consisting of a few individuals in creeks and banks populated by larger numbers of conspecifics. These fish will readily interact when exposed to another individual and the nature of this interaction depends on dominance status, familiarity, physiological state, etc. As a nocturnal species, *A. leptorhynchus* tend to hide during the day and are territorial regarding hiding places. They also prefer confined spaces, although they do explore their environment [Moller, 1995].

Communication signals and behavior have been particularly well studied behaviorally and physiologically. Social interactions in weakly electric fish are largely mediated by their electrosense and the electric field they generate. *A leptorhynchus* continuously generate a weak (mV) electric field that oscillates in polarity at a frequency of 600 to 1000 Hz. The base frequency of this electric organ discharge (EOD) can signal sex, maturity, and individual identification because EOD frequency is stable in an individual [Harvey-Girard et al., 2010; Zakon, and Smith, 2009].

When the electric fields of two nearby fish interact, the resulting perceived electrical signal has an ongoing modulation in amplitude called a beat. The frequency of this beat allows the fish to estimate the EOD frequency of a conspecific, relative to his own and remains as an ongoing signal present in the background of any conspecific interactions [Hopkins, and Popper, 2005]. These fish produce transient EOD modulations[Turner et al., 2007] during social interactions.

One such modulation is the well characterized jamming avoidance response (JAR). The JAR is elicited when two fish with relatively similar frequencies come in close proximity. The two fish change their baseline frequency by a few Hz thereby avoiding a slow beat modulation that may jam their ability to electrolocate [Heiligenberg, 1991]. The main communication signals are very short and are characterized as transient increases in EOD frequency, known as chirps, which usually last 10 to 100 ms [Turner et al., 2007]. Transient EOD modulations like chirps and JAR are key components of conspecific interactions including aggressive and agnostic interactions or courtship [Henninger, 2015].

The presence of CB_1_ receptor in the brain of *A. leptorhynchus* has been confirmed [Harvey-Girard et al., 2013] and its distribution in the nervous system presents many similarities with other teleost fish or with mammals. Nothing is known about the role of the CB system in gymnotiformes’ physiology and behavior. Our general goal with the present study was to make a first step towards using this powerful model system to better understand the role of the cannabinoid system in shaping the neural substrate of behavior. In doing so, our behavioral analysis reveals an intricate effect of CBs on social interaction. We first characterized the overall behavioral effect of the CB receptor agonist Δ^9^-tetrahydrocannabinol (THC) on *A. leptorhynchus* behavior. The initial results allowed us to hypothesize that THC-induced behavioral alterations were context dependent.

## Methods

### Fish care and use

Adult male and female wild caught *Apteronotus leptorhynchus* were acquired from commercial suppliers. Typically, fish were housed in 100 L tanks containing up to 10 individuals. PVC tubes were placed in the tank to provide hiding spots and environmental enrichment. Water condition was monitored daily and maintained in normal ranges (e.g., pH 6.5-7.5; conductivity 200-300 μS; temperature 26.5-27.5 °C). Water conditions in experimental tanks were identical to home tanks. Fish were acclimated in the laboratory at least three weeks before being used for experiments. Fish from a given tank were kept together, and changing the tank composition was avoided. Housing facilities were on an inverted 12:12 light cycle; experiments were conducted during the fish’s active nocturnal phase. Fish used for these experiments were identified with a label on the tank and, when needed, fish were individually marked with small fluorescent elastomer implants under the skin (VIE tags; Northwest Marine Technologies Inc., Shaw Island WA). We thereby ensured that a fish was not used for experiments until it had recovered for at least 7 days, and that no behavioral, electromotor, or physical symptoms remained from the previous experiments. Fish care procedures and the experimental procedures described below were approved by WVU’s Animal Care and Use Committee.

### THC injections

Studies on the behavioral effects of systemic THC injection in other species helped us delineate the appropriate dose range and testing condition to reveal the most likely effects. Three concentrations were tested spanning a range of 33 mg/kg to 100 mg/kg, which is known in other species to reach levels considered high but non-lethal [Varvel et al., 2005]

Fish were individually weighed prior to injection. Injection volume was 0.01 ml/g. THC was generously provided by the NIDA Drug Supply Program (Bethesda, MD) and dissolved in a vehicle consisting of 5% ethanol, 5% Kolliphor EL (Sigma-Aldrich, St. Louis, MO), and 90% saline [Kinsey, and Cole, 2013]. THC concentration in the solution was 0 mg/ml (control), 3.3 mg/ml, 6.7 mg/ml, or 10 mg/ml, leading to injections of 0, 33, 67 or 100 mg/kg of body mass. Single use 0.3 ml syringes with 30-gauge needles were used for intramuscular injection in the epaxial muscle. These intramuscular injections were very rapid with minimal disturbance to the fish. Long-term visible injuries to the skin or muscle did not occur.

### Chirp Chamber experiments

*A. leptorhynchus* spend a large portion of their time in small burrows and root masses where they find protection from predators. In laboratory aquaria, fish are given PVC tubes for use as hiding places. We modified one of these PVC tubes by opening the sides and replacing them with plastic mesh in order to be able to stimulate and record from the fish while they were in the tube. *A. leptorhynchus* individuals readily responds to conspecific communication signals when hiding in the tube and this paradigm is widely used to quantify the responsiveness of the fish to various stimuli [Harvey-Girard et al., 2010].

This experimental setup consisted of a 75 L aquarium containing a chirp chamber (i.e. the modified PVC tube). A stimulation dipole was placed 10 cm apart on one side, 20 cm away from the chirp chamber, and a recording dipole was attached to the extremities of the tube. The stimulation and recording dipoles were in perpendicular orientation so that the recording dipole picked up the fish’s signal strongly, but not the stimulus we delivered.

Individual fish were introduced in the chirp chamber, and the opening was closed to prevent the fish from leaving the tube. After 5 min acclimatization, the first set of stimuli were played and electromotor behavior was recorded. The fish was then taken out of the tank, injected, placed back in the tank, and after 15-20 min, the fish was guided into the chirp chamber. After an additional 5 min, the set of stimuli was repeated. Stimuli consisted of sinusoidal signals mimicking a conspecific EOD. The frequency was adjusted relative to the fish’s own frequency to create beat amplitude modulations of specific frequencies ranging from −120 to +120 Hz. To do so, the fish’s EOD frequency was measured before each stimulation using the frequency estimate of the fish’s signal displayed by the oscilloscope (GwInstek GDS-2074A, Good Will Instrument Co., Taiwan). Some stimuli were simply constant EOD frequencies whereas others contained transient chirp modulations. These modulations were Gaussian increases in frequency of 120 Hz that lasted 14 ms and were presented twice per second, which corresponds to type 2 chirps used in numerous previous studies [e.g. Marsat et al., 2009]. Each stimulus type was played for one minute with 2-minutes pauses in between. Stimuli were created and played with custom Matlab (Mathworks, Natick, MA) scripts controlling the computer’s generic sound card. Signals were produced at 44100 Hz and isolated before being sent to the tank (model 2200 Analog stimulus isolator, AM Systems, Sequim, WA). Stimulus intensity was calibrated to cause EOD intensities similar to that produced by an average conspecific. To achieve this calibration, we placed a small recording dipole in the chirp chamber perpendicular to the chamber’s main axis. We recorded and averaged the intensity of the signal emitted by 10 fish placed directly above the chirp chamber (perpendicular to its axis). The stimuli where adjusted to cause the same intensity. Signals were amplified (model 1700 amplifier, AM Systems) and recorded (sampling rate of 44100 Hz) with the same sound card and custom Matlab script used for stimulation. Analysis of the recordings was also done using Matlab scripts by visualizing the recordings as spectrograms (see Fig 2). Changes in baseline EOD frequency of the fish (JAR response) and the occurrence of chirps were visually identified by the experimenter (blind to experimental conditions) and recorded through the graphical interface.

**Figure 1:**
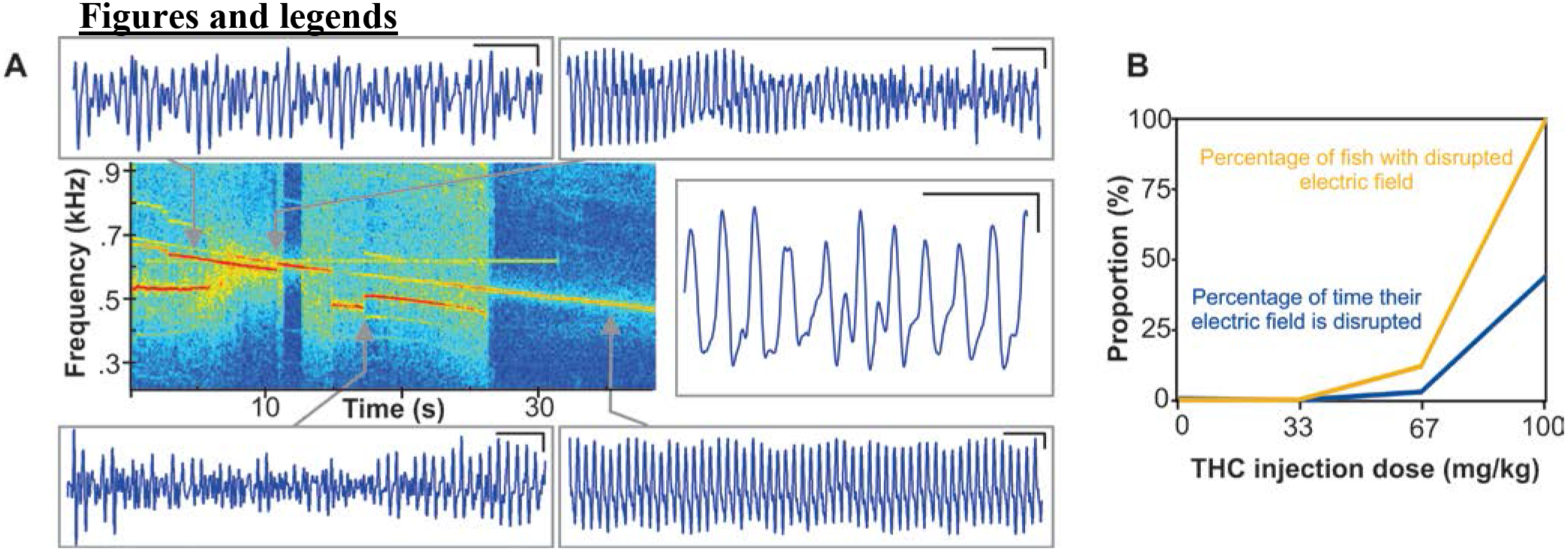
Disruption of the EOD at high dose of THC. **A.** Spectrogram of the disrupted EOD of a single fish with insets of EOD waveform excerpts. The excerpt (i.e. time 0 on the spectrogram) starts 10 min after injection. The main EOD frequency at any given moment can be seen as the dark red-brown line. A normal EOD would be visualized on the spectrogram as a single flat line. Here the EOD frequency decreases, fluctuates, has multiple peaks of frequency power, and is overall very unstable. Four insets (top and bottom) taken at different points on the spectrogram show the details of the time varying EOD waveform, with a fifth inset (right) providing a more detailed view of an example EOD being unstable. In the insets, black scale bars in the upper right corner of each inset represent 10 ms and 1 mV. **B.** Propensity of EOD disruption as a function of THC dose. We quantified both the proportion of fish that displayed any disruption and the proportion of the time the EOD is affected in these fish during the 20 min following injection (n=8 for each dose <100 mg/kg and n=4 for 100mg/kg).

**Figure 2:**
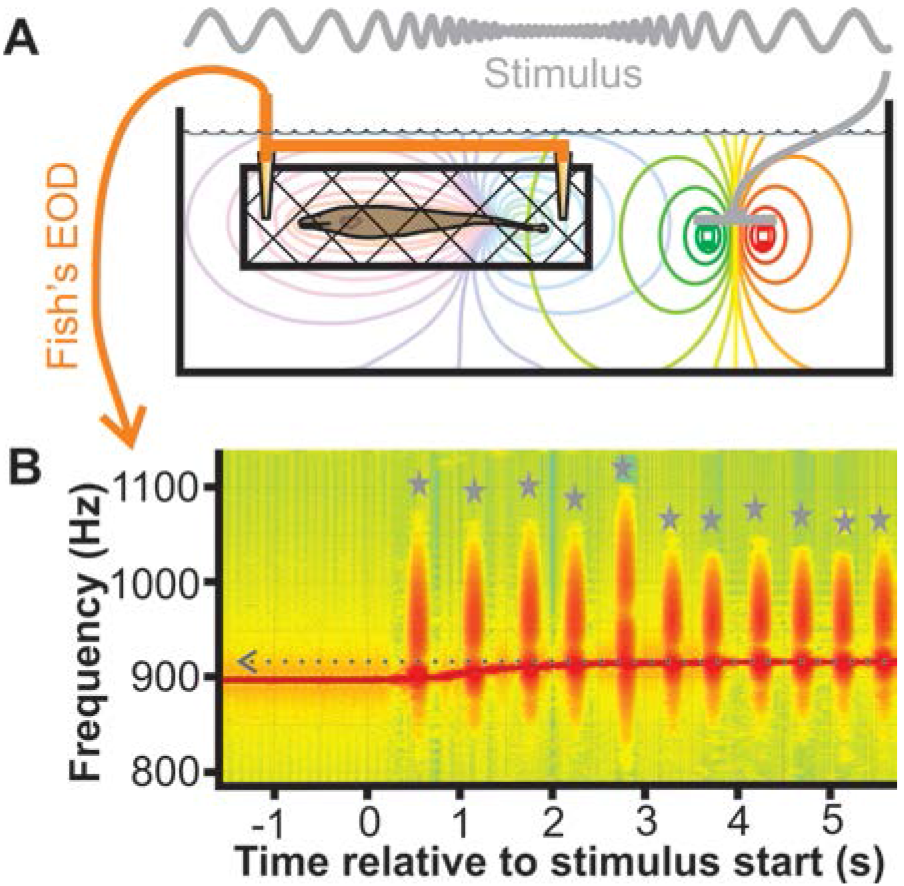
Chirp Chamber stimulation and recording visualization. **A.** Illustration of the recording set-up. The fish is placed in a mesh tube wired with a recording dipole to capture the fish’s EOD (depicted by purple shades electric field lines). A stimulation dipole is placed on one side to mimic a nearby conspecific (red-green dipole and field lines). **B.** Recordings are displayed as spectrogram (showing intensity as a function of frequency and time). In this example (stimulus frequency: beat of −10 Hz) chirps are highlighted with grey stars and the increase in baseline frequency (JAR) with the grey arrow.

### Aggressive encounter experiments

Fish were injected and placed in a small 30 cm × 30 cm tank with 10 cm water depth in order to constrain movements of the fish mostly to 2D. The tank was further divided in half with a diagonal, plastic mesh panel. The half used for experiments contained only the stimulus and recording dipoles, whereas the opposite half contained the inlets/outlets for warm water and an air bubbler. The recording dipole was placed at opposite corners of the tank, and the stimulus dipole (5 cm between poles) placed in the middle of the compartment, perpendicular to the recording dipole. After a variable acclimatization time (0.5, 1, 2 or 4 hours depending on test group; see results), ten one-minute-long stimuli were presented with 2 min pauses in between. The stimuli consisted of an EOD with a frequency that was 10 Hz below the fish’s own, and small chirps identical to the ones presented in the chirp chamber experiment. The stimulation and recording equipment were also identical to the chirp chamber rig. The tank was enclosed in an opaque box to block all external light. Infrared LED illumination was provided from below and video recordings were captured using a webcam (model C920, Logitech, Newark, CA) with its IR filter removed. In a first set of trials (waiting time of 20 min). Video recordings were scored by visually identifying and counting lunges. Lunges were operationally defined as rapid forward movements. These stereotyped motor patterns are easily identifiable. In a small subset of recordings, a second experimenter re-scored the trial. In every case the lunges identified were identical between the first and second experimenter (r = 1). For the second set of experiments, lunges were identified [Hupé and Lewis., 2008] and EOD recordings were processed as described in the previous set of trials, following quantifying chirping and JAR frequency changes. In addition, the videos were processed with ANY-maze video analysis software (Stoelting Co., Wood Dale, IL) to track fish position. The software identified the fish based on contrast and used the tip of the frontal end to determine the x and y coordinates for each frame. We verified visually that the program correctly identified the fish and its frontal edge. Using the coordinates, we quantified several aspects of the fish’s locomotor behavior, including time near the stimulus dipole (i.e., ≤ 2.5 cm from either pole), distance traveled, and mean swimming speed. These different measures gave similar results, with swimming speed showing less variability, and we therefore show swimming speeds in the Results section.

### Social setting experiments

This set of experiments used a large 1 m × 1 m tank (20 cm water depth) divided into 8 compartments, with a closed, ninth central compartment. These compartments were connected to each other by 3 cm wide openings starting at 5 cm from the bottom of the tank and extending up to the surface of the water. Each compartment contained a hiding tube, carbon rod recording dipoles located in opposite corners, and inlets and outlets connected to a sump for circulation of clean, warmed, and oxygenated water. The tank was covered with an opaque box and kept in the dark during the recordings. An IR video camera was used to monitor the fish without disrupting the experiment. After being injected with THC or vehicle, four fish were placed in different compartments on opposite corners of the tank. The fish in the “familiar environment” group were left to acclimatize to the tank for 24 hr before injection. EOD signals from each compartment were recorded for 3 hrs after injection. Signals were amplified (model 1700 amplifier, AM Systems), and data were acquired at a sampling rate of 20 kHz for each of the 8 channels using an AD/DA board (PCIe-6343, National Instruments, Austin, TX). Analysis relied on a custom Matlab script to visualize the recordings as spectrograms, as described for the chirp-chamber experiments extended to accommodate 8 separate recording channels. For each 1 s of recording, an experimenter determined where each fish was by comparing the EOD frequencies and amplitude present in each compartment. Because the opening between compartments was small and the recording dipole orientation was kept constant in each compartment, the strength of an EOD signal of a given fish was always stronger in the compartment that contained the fish compared to the recordings of this signal from adjacent compartments. Simple comparison of the spectrograms and power spectra of the recordings from each compartment allowed determination of where the signal was strongest for a given fish’s EOD frequency. In a separate set pilot of experiments, results of 120 single estimates of fish position per this method, were correlated with the position of fish in a video recording of the tank to obtain a 100% match. Rather than tracking the position of individual fish, the number of fish present in each compartment at the beginning of each 1 s portion of recordings was quantified. Movements of fish from one tank to the next were reflected as a change in the number of fish in the corresponding compartment. The variability of the number of fish per compartment (i.e., “position variability”) was used as a measure of fish movement between compartments. In very rare cases, if two fish traded respective compartments within 1 s, the fish count in these two compartments would not change. In such rare cases our estimated fish measure may slightly underestimate fish movement across compartments. Given that such an underestimation, however small in magnitude, would occur in equal proportion across chambers and test conditions, risk of type 2 error is minimal. Prior to statistical analysis, we ascertained that our data were normally distributed with Matlab, using a Kolmogorov-Smirnov test. Normality was confirmed in all cases except for the results displayed in figure 3, and thus parametric or non-parametric statistical tests were applied accordingly.

## Results

### Initial observations

Five different behavioral paradigms were used as detailed in the coming sections. The first one was simple observation and electrical monitoring of fish that were injected, released in a 75 L tank, and free to move. The fish were monitored for 20 min as an acclimatization step before the chirp chamber recordings described in the next section. At high doses, the behavioral effects were obvious and rapid, starting within minutes of injection. At the highest dose of 100 mg/kg, nearly every fish would display the same behavior. Once released in the tank post injection, fish would swim rapidly around the tank once or more, then drop to the bottom of the tank and become nearly motionless. They would occasionally initiate a short bout of energetic swimming that would last a few seconds. The period of immobility lasted up to several hours. Lower doses had more moderate effects on swimming in a dose-dependent manner and higher doses had more severe effects (see next sections).

The most surprising effect of the high dose (i.e., 100 mg/kg) injection was on the generation of the electric field. It is important to point out that these fish generate an electric field continuously throughout their adult lives. Drastic procedures, such as spinal transection, must be performed to prevent the generation of the electric field. However, high doses of THC disrupted and often eliminated the EOD. The typical initial effect was for the EOD to be erratic, decreasing in frequency with time, then suddenly jumping back up with bouts where the EOD had multiple frequency peaks (Fig 1A). In the most affected fish, the EOD generation stopped and started abruptly with longer and longer periods of silence, so that the EOD was silenced within 10-20 min. We would like to point out as a casual observation that, although all other observable behavioral effects remitted within hours, fish that had their EOD disrupted and silenced did not recover it until days later. This recovery seemed to be slow with the EOD gaining amplitude with time. We did not quantify this long-term phenomenon, which should be the topic of a thorough investigation of the effects of CB agonists on the EOD generating mechanisms. We did, however, quantify the propensity for this initial disruption in EOD generation. The disruption was either clearly present or not at a given time, with no graded transition between states. We quantified the percentage of fish that displayed these erratic EOD disruptions and the percentage of time the disruptions were observed (Fig 1B). The effect was observed in all the fish for injections of 100 mg/kg THC. Lower doses did not cause this effect except for a single trial where a fish injected with 67 mg/kg THC displayed a brief interruption.

### Electromotor response to communication signals in a hiding chamber

The first set of stimuli tested consisted of simple beat modulations that replicate the sinusoidal AM experienced by the fish when in close proximity to a conspecific. Fish were tested before (Fig 3A) and after THC injection (Fig 3B). The typical JAR was not systematically affected by THC, and there was no effect of dose on the JAR (Kruskal-Wallis, p=0.4). This lack of effect of THC on the JAR was useful in confirming that effects described in the following sections do not simply reflect generalized lack of responsiveness of the fish, but rather selective effects of THC on some behaviors but not others.

**Figure 3:**
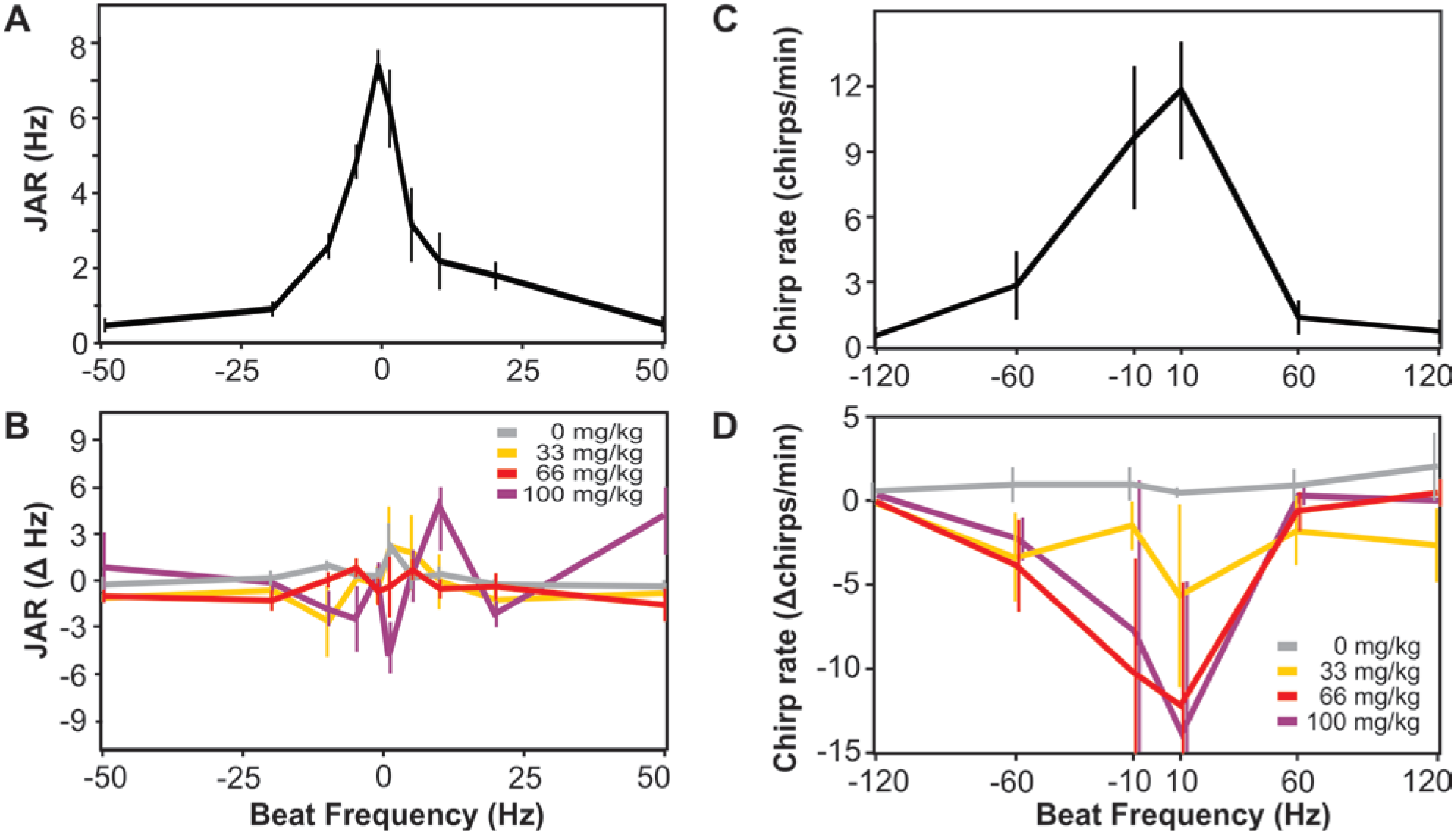
Selective effect of THC on responses to communication signals. **A.** Baseline JAR response prior to THC injection as a function of stimulus frequency. Stimuli were simple beats (sinusoidal AM) created by presenting an artificial EOD at the indicated frequency above or below the test fish’s frequency. The average (± s.e., n=28) across the fish used in B is shown here. **B.** Change in JAR response displayed in A after THC injections (post-pre) at different doses (average ± s.e., n=8 for each dose <100 mg/kg and n=4 for 100mg/kg). **C.** Chirping rate in response to chirp stimuli of different beat frequencies. The average (± s.e., n=28) across the fish used in D is shown here. **D.** Change in chirping response displayed in C after THC injections (post-pre) at different doses (average ± s.e., n=8 for each dose <100 mg/kg and n=4 for 100mg/kg).

Chirp stimuli were also presented and consisted of constant beats (frequencies of −120 Hz to 120 Hz) interrupted twice per second by a small chirp (type 2). The fish tended to respond to these signals with chirps, often produced in an echo response, and it is known that low frequency beats elicit more response chirps than high frequency beats (Fig 3C; [Hupé, and Lewis, 2008]). After THC treatment (Fig 3D), chirping response was significantly affected in a dose-dependent manner (Kruskal-Wallis, p=0.016). Chirping was completely suppressed at the two higher doses, reduced at the weaker dose (33 mg/kg), and unaffected for vehicle injections. Chirp-chambers constitute a safe, low-stress environment, and our results show that in such conditions the fish’s tendency to chirp in response to a conspecific signal is reduced by a CB agonist.

### Aggressive encounter scenario

One of the most frequent and obvious social behaviors of these fish is agonistic interactions. Agonistic behavior can be easily triggered by putting two fish in a confined space [Hupé and Lewis, 2008]. Similarly, agonistic behaviors can be artificially elicited in one fish presented with a mimic of the signal from another fish. The probability of observing aggressive interactions can be increased by (1) removing any hiding place, and (2) using a stimulus EOD that is close in frequency to the fish (i.e., low frequency beat) and that contains chirps. We thus constrained a fish to one half of a 30 cm × 30 cm tank, in the middle of which a stimulation dipole was placed (Fig 4A). After a 20 min acclimatization, the stimulus was presented for 1 min followed by 2 min of rest without the stimulus. This stimulus-rest cycle was repeated 10 times.

**Figure 4:**
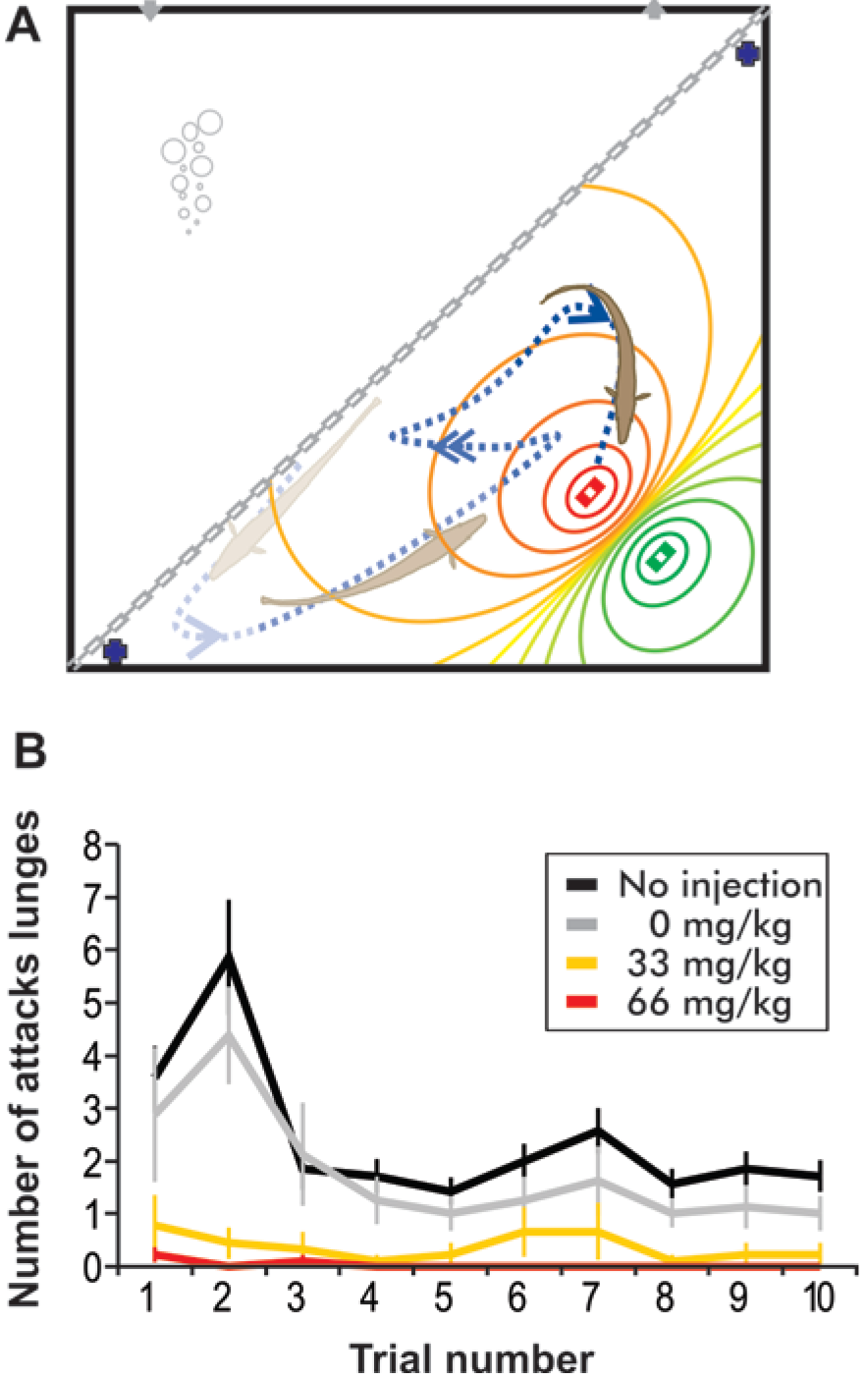
THC decreases attack behavior in agonistic situations. **A.** Agonistic situations are created by placing the fish in a confined space. Here only half the square tank is used, and access to the other half containing an air bubbler and water recycling inputs/outputs is restricted with a mesh barrier. A stimulation electrode is placed in the middle of the test compartment (red-green dipole creating an electric field) and recording electrode at each extremity (black circles). Movement of the fish is recorded with an IR camera while a stimulus with a low frequency beat and chirps is played. **B.** Decrease in attack lunges with increased THC dose. For each of the 10 bouts of stimulation during an experiment, the number of attack lunges produced by the fish is quantified. The mean (± s.e.) is displayed for experiment sets with different THC treatments (n=8 each for no injection and 33mg/kg; n=10 for 0 mg/kg and 66 mg/kg).

As expected, the fish responded to the stimulus with characteristic aggressive swimming movements consisting of high speed movement towards and around the stimulation electrodes and stereotypical lunges [Hupe et al., 2008]. *A. leptorhynchus* fight by lunging and biting at each other. Similarly, when presented with an electrical mimic, the target fish lunged toward the stimulus dipole and made biting motions in the target areas near the probe that corresponded with the body position of a real fish. These lunges were most frequent in the first few trials of sequences of 10 stimulations (Fig 4B), indicating that the fish habituated to the artificial fish stimuli. Injections of weak and moderate doses of THC significantly decreased the propensity of the fish to lunge (ANOVA, p=0.02). Note that this suppression of lunging was not due to the physical injection procedure, because the responses of vehicle control fish did not differ from non-injected controls (ANOVA, p = 0.3).

We used this small-tank agonistic stimulation paradigm to evaluate the duration and time course of the behavioral effects of THC. 67 mg/kg THC dose was used in the following experiments based on the data presented above that 67 mg/kg THC had clear behavioral effects without the risk of the fish losing its EOD and movement ability seen at 100 mg/kg. We repeated the aggressive encounter experiment described above but varied the latency between the injection and the stimulation (from 0.5 hr to 4 hr). In addition to lunging, we quantified each fish’s JAR response, average swimming speed, and chirping response. JAR was unaffected by THC, confirming once again that the effects observed are selective (Fig 5A; ANOVA, p=0.18). The swimming speed was decreased by THC in the first 2 hrs following injection (Fig 5B; ANOVA, p= 4.10^−8^). This decrease diminished after 2 hrs and vanished by 4 hrs (Tukey HSD, p<0.01 for 0.5 to 2 hrs and p=0.32 for 4 hrs). We suggest that this time course represents the time course of the overall effect of THC injections and thus argue that for these type of injections, THC will affect the fish’s behavior for 2-3 hrs.

As shown in figure 4B, THC injected fish lunged significantly less frequently than controls when tested within 1 hr of injection (Fig 5C; ANOVA, p=7.10^−6^; Tukey HSD, p<0.05 for 0.5 and 1 hr and p>0.05 for 2 and 4 hrs). When control fish were newly place in a tank and presented with chirp stimuli, they respond with a relatively high amount of attack movements; this propensity decreases if they have been acclimated to the tank for several hours. THC injections reduced the frequency of these initial aggressive movements to the same level as control fish that had been acclimated. This data shows that THC decreased aggressive lunges in fish placed in an aggressive encounter scenario.

**Figure 5:**
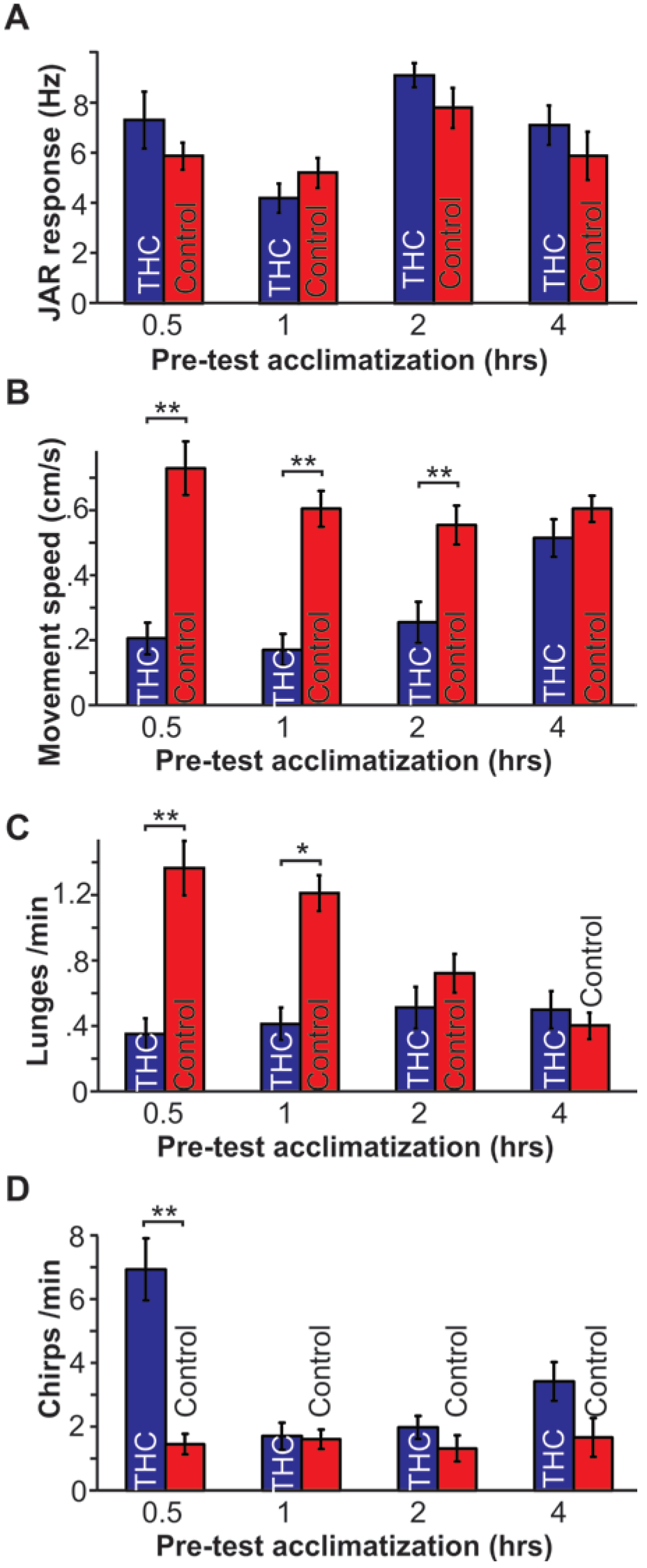
THC selectively affect movement and social behaviors in the first hour post injection. Four aspects of the behavior (means ± s.e.; n=8 for each group) during the agonistic situation described in Fig. 4 are quantified during about stimulation performed after a variable acclimatization period following THC injection. **A.** JAR response (as illustrated in Fig, 2). **B.** Average movement speed during stimulation. **C.** Attack lunges produced by the fish. **D.** Chirps emitted by the fish in response to the stimulation. Stars indicate statistically significant differences (**: p>0.01; *: p>0.05)

In addition to lunging, THC also significantly increased chirping responses in this context but only when the fish was provided a short time to acclimate to the new environment (Fig 5D; ANOVA, p= 0.003; Tukey HSD, p>0.05 for 1, 2, and 4 hrs). Interestingly, the THC-induced increase in chirping observed here is opposite to the effect observed in the chirp-chamber. We will discuss the interpretation of this result in the discussion, but it is noteworthy that previous research indicates that chirping can be either a sign of aggression or a signal to de-escalate aggression when chirping is produced as an echo response between the two fish [Hupé, 2012]. We therefore calculated the probability that the chirps produced were in response to one of the stimulus chirps (i.e., following it closely in time). Focusing on the experiments where the fish was tested 0.5 hr after injection, we calculated that 65% of the chirps produced by the fish are indeed echo responses. The likelihood that chirps produced are echo chirps (i.e., chirps preceded by stimulus chirps) is similar for THC-injected vs. control fish (0.61 ± 0.21 vs. 0.7 ± 0.18 respectively; T-test, p=0.37). Considering that echo chirps are thought to be signals to de-escalate aggressiveness, the increase in chirping is consistent with the decrease in lunging and aggressiveness.

### Social environment in novel versus familiar conditions

The results of our experiments so far indicate THC affects *A. leptorhynchus* social behavior in a context-specific manner. For example, in the chirp chamber, the fish decreased communication behavior, whereas in the freely-swimming aggressive scenario, the fish increased communication behavior, potentially to de-escalate aggression. We next aimed to determine if these different effects on social interaction were indeed the consequence of the two different contexts. To do so, we used a large tank that was separated into small interconnected compartments, each containing a single hiding tube for the fish (Fig 6A). This arrangement was designed to provide a comfortable living space where several fish can reside without stress caused by limited space or hiding tubes. The experiment involved 4 fish being placed in the tank. The “familiar” scenario used 4 fish from the same housing tank, with an established dominance hierarchy. The fish were placed in the tank 24 hrs before the start of the experiment. Thus, the socio-physical environment was also familiar at the beginning of the experiment. The “novel” scenario used 4 fish from 4 different home tanks. In addition, there was no acclimatization to the test tank before testing began. These fish were therefore subject to a novel socio-physical environment. Data were collected over the 3 hrs following injection (see methods) of either 67 mg/kg THC or vehicle. By visualizing a spectrogram of the EOD signals recorded from each compartment, fish location chirping rates were easily recorded (Fig 6B).

**Figure 6:**
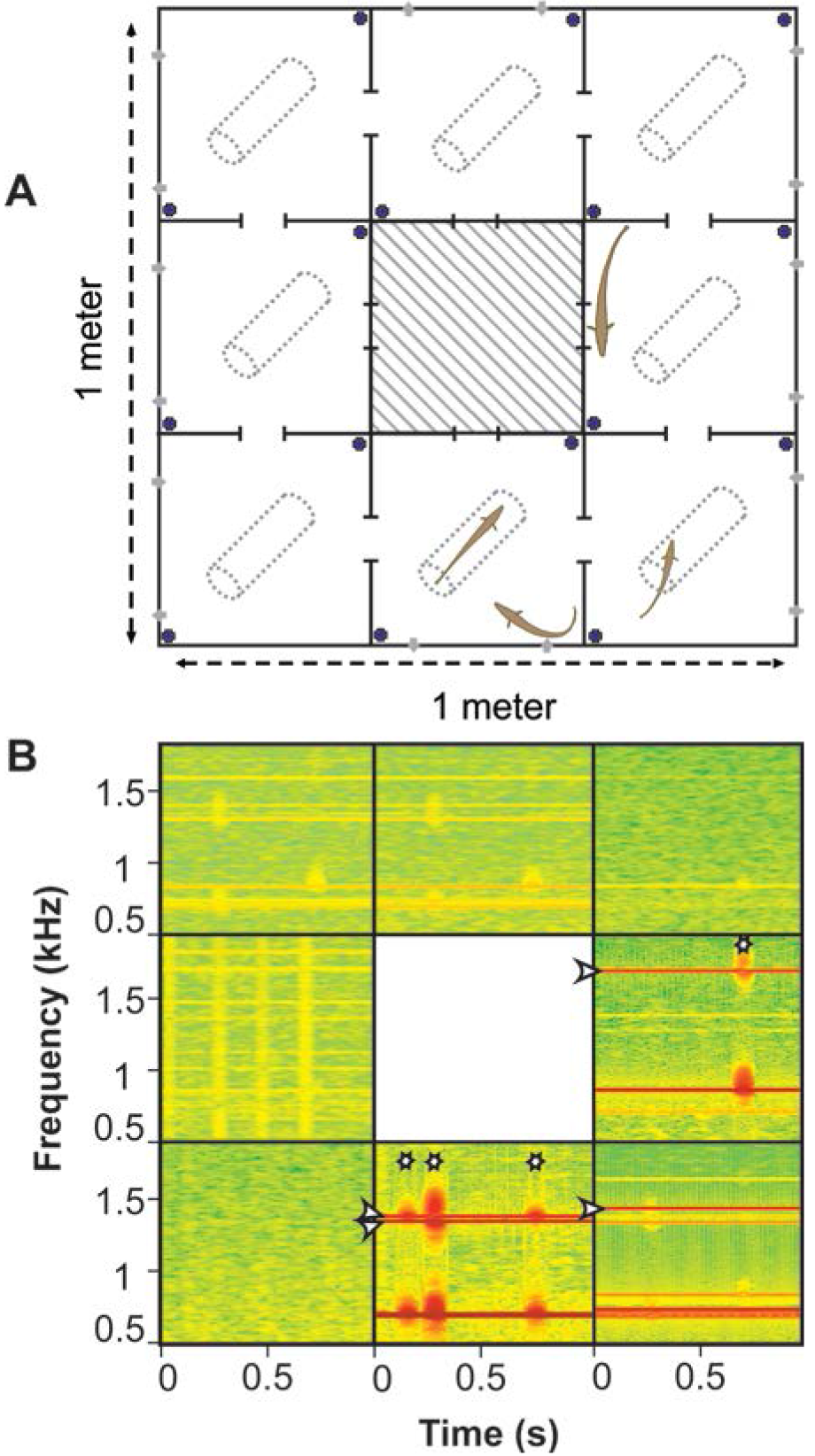
Characterizing the effect of THC on social interactions in a small group of fish. **A.** A large compartmentalized tank is used to allow the fish to explore and cluster in individual compartments. Each compartment contains a hiding tube and is connected to others by small openings. A pair of recording electrodes (black circles) is placed in each compartment which are also equipped with a water input/output (grey dots) providing cleaned, warmed and oxygenated water. Four fish are initially placed in the four corners of the tank. A 3 hr recording session is started after injection. Two types of trials were performed: in the novel environment trials, fish were placed in the tank immediately after injection and with unknown tankmates whereas, in the familiar environment trials, fish used are long-term tankmates and are placed in the testing tank 24 hr prior to injection. **B.** Recordings from each compartment were displayed as spectrograms (NB: when needed to resolve ambiguities, we also looked at power spectra where the time dimension is collapsed). Fish EODs show up as dark lines in the 0.6-1 kHz range and the 1^st^ harmonic of their EOD in the 1.2-2 kHz range. Based either on the baseline frequency or 1^st^ harmonic trace, the position of the 4 fish at the beginning of each 1 s time frame was determined (white arrow) and chirps where marked (white star) to be counted.

Consistent with the decreased motor activity observed in the other tests, the THC-injected fish changed compartments less often than their control counterparts (Fig 7A; ANOVA followed by Tukey HSD, p= 6.10^−7^ for the effect of THC). This effect was most pronounced shortly after injection (Fig 7B) and dissipated 2.5 hrs after injection. Novel vs familiar treatment groups showed no differences in position variability (ANOVA followed by Tukey HSD, p= 0.38 for the effect of socio-physical environment).

**Figure 7:**
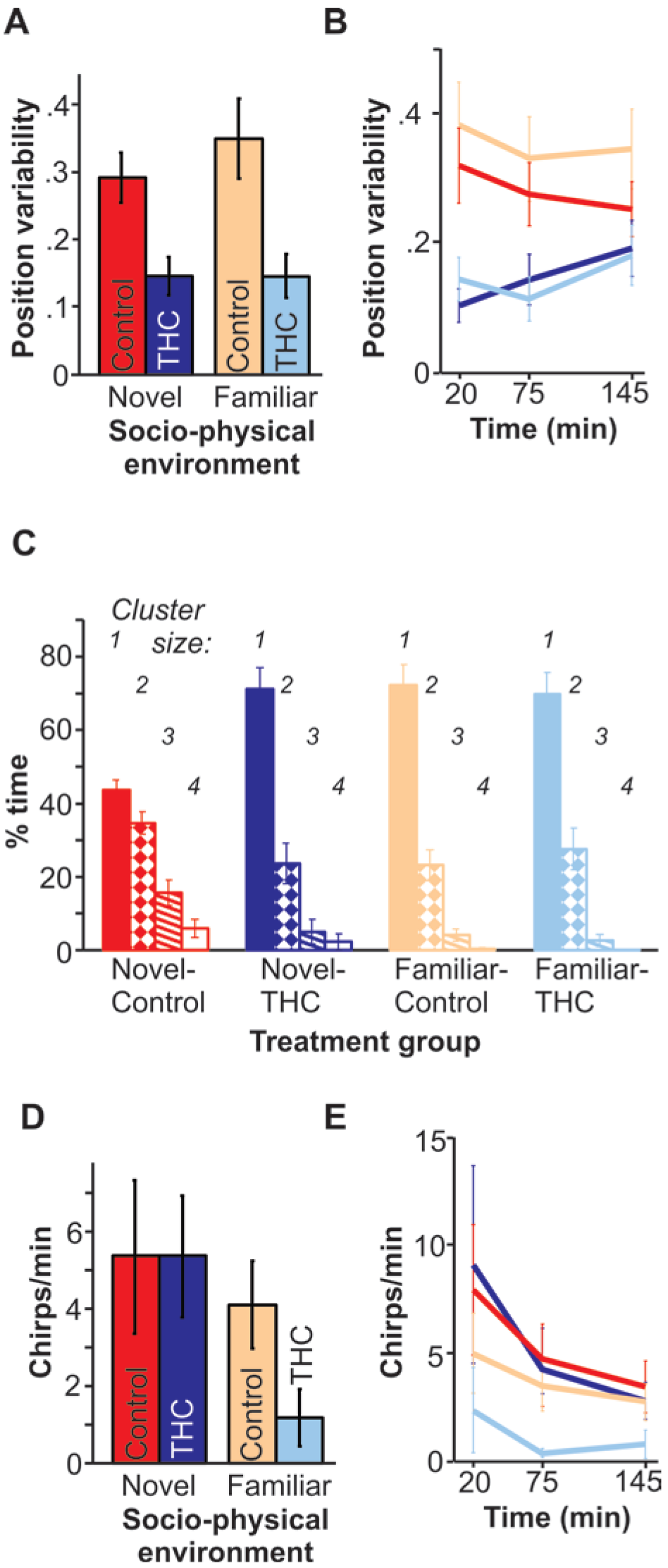
THC has different effects on fish in familiar versus novel socio-physical environments. Using the experimental paradigm described in Fig.6, we quantified several aspects of the fish’s behavior at different times during the 3 hr trial (i.e., 30 min windows starting at 20 min post injection, 75 min or 145 min). Ten experiments (single 4 fish) were performed for each treatment group (novel/ familiar, THC/control). Averages across fish (± s.e.; n=24 fish) are shown either as an average across the three analysis windows (A,C,D) or separately (B,E). **A,B.** Position variability decreases with THC injection independently of the socio-physical environment conditions. Position variability was quantified via the change in composition of each compartment (see methods) and reflects the tendency to move and explore the environment. **C.** Clustering is decreased by THC but only in novel environments. The proportion of time spend in a compartment by itself (cluster size 1; filled bars) or with other fish (clusters 2-4; patterned of empty bars) is displayed for the different treatment groups. **D, E.** Chirping is decreased by THC but only in familiar environments. Chirp rate per individual fish is displayed for each treatment group.

One of the most striking effects of THC injection was observed for the novel condition, by comparing the dispersion of fish throughout the compartments (ANOVA p=5.10^−6^ comparing only the % time found in “clusters of 1”). Control fish were very likely to be found in groups of 2-3 individuals, often even sharing the same tube. This huddling behavior is observed in a variety of fish species in response to stressful situations such as novel environments [Johnston, and Glasgow, 2015; Miller, 1963]. Although no publication documents this behavior in gymnotiformes, we routinely observe this clustering when a group of fish is placed in a novel tank. It was therefore not surprising to see a higher level of clustering in the novel-control test groups compared to the familiar-control test groups (Fig 7C). In the present study, clustering in the novel-THC group was as low as clustering for fish in a familiar environment. Clustering was also low in the familiar environment, and THC-injection did not further lower clustering.

THC injections did influence chirping propensity: the familiar-THC groups chirped less than their control counterparts (Fig 7D; T-test for familiar control vs THC, p=0.013) throughout the duration of the recording (Fig 7E). Both groups exposed to the novel environment chirped at similarly high rates (T-test, p=0.69). THC injection thus affected distinct aspects of the behavior in novel vs. familiar environments. In novel environments, they clustered less than control groups but continued communicating with each other at the same rate whereas THC-injections decreased chirping in familiar environments.

## Discussion

The behavioral effects of cannabinoid agonists described here are overall consistent with observations in other species in which one of the most marked effects is a decrease in locomotor activity ([Rodriguez de Fonseca et al., 1998]; e.g., Figs 5B &7A). The influence of THC on locomotor activity in *A. leptorhynchus* led to a decrease in swimming and at very high doses, THC injections resulted in a disruption of the electromotor activity. Our study focused on communication behavior and social interactions. Our data indicate that THC injection did not merely induce a general decrease in all behavior, as would be expected if the animals were simply sedated. Rather, we see specific effects on some behaviors and lack of effects on others (i.e. JAR). Chirping in THC-injected fish was decreased in chirp-chambers experiments when presented with simple beat stimuli. It also decreased when placed in familiar context with several other fish but chirping was not affected by THC in unfamiliar context. In contrast, THC caused an increase in chirping in isolated freely moving fish presented with chirp stimuli. Our experiment also showed that THC injections could reduce certain social behaviors: it decreased aggressive lunges in fish placed in a new tank and presented with chirp stimuli, and decreased the likelihood of fish to cluster in groups when placed in a new social and physical environment.

CB1 receptors are found in a wide range of regions in the brain [Harvey-Girard et al., 2013] including motor and sensory areas as well as a high concentration in higher brain areas (forebrain). Our study clearly shows that electromotor activity is disrupted at high THC doses. Harvey-Girard et. al. [2013] showed that the pre-pacemaker nucleus, a region modulating the pacemaker that drives the generation of the electric field, is rich in CB receptors. We cannot rule out the possibility that the EOD generation is disrupted at another site than the pre-pacemaker nucleus. Nevertheless, lesions in the prepacemaker nucleus can cause changes in EOD frequency and regularity [Moortgat et al., 1998]. Thus, it seems plausible that CBs could affect the pre-pacemaker dynamic leading to observed changes in EOD output.

It could be suggested that most of the effects observed in this study result from the influence of the CB system on the motor system. For example, it could be suggested that a generalized depression of the motor system in THC-injected fish causes the slower swimming speed, the decrease in aggressive lunging or the decreased clustering. We argue however that an effect on the motor system is unlikely to explain all of our observations. Specifically, we show that THC sometimes increases chirping and sometimes decreases chirping. Also, clustering is affected in a context-dependent manner (Fig 7C) whereas the effects on the amount of movement were not context-dependent (Fig 7A). We therefore suggest that although an effect on the motor system can explain some of our observation, other effects take root somewhere else.

The lower level of the sensory system involved in electrocommunication is also a major focus of research in weakly electric fish. In particular, the ELL and the feedback pathway through parallel fibers have a well understood role in processing the electro-sensory signals (for reviews on the topic, see [Hopkins, and Popper, 2005; Maler, 2007; Marsat et al., 2012]). This network has several features specialized for processing communication signals. The main output cells of the ELL, the pyramidal cells, receive massive feedback inputs from higher levels to enhance the processing of communication signals. One of these sources of feedback, the granule cells of the caudal cerebellum, express CB receptors, and involves plasticity [Bol et al., 2011]. CB system may thus influence the processing of communication signals as early as the primary electrosensory area of the CNS. CB1 receptors are also present in higher areas of the sensory system such as the torus semicircularis [Harvey-Girard et al., 2013]. Once again however, it seems unlikely that a change in sensory processing play a large role in explaining the main behavioral effect of THC observed in this study. For example, chirping decreased in the chirp-chamber experiment eventhough the JAR was unaffected. If THC cause a change in responses to beat stimuli in this context, you would expect both behavior to be affected. Also, chirping does not always decrease in response to THC injection, a change in perceptual abilities would be expected to affect responsiveness in all situations. Nevertheless, we do not rule out the possibility that theeffect of CB on the sensory system could have affected the behavior of the fish in our experiments.

It is likely that whole-brain dynamics, including higher-brain areas, is affected by the CB system. A possible explanation for the context-dependency of some of the behavioral effects of THC injection is that different contexts cause state changes in the animal and influence levels of stress, anxiety or aggressiveness. We detail how our results support this suggestion in the following two paragraphs.

THC caused a decrease in clustering in a novel social and physical environment but not a familiar one. This change in clustering can be understood based on the typical reaction of these fish to novel environments. Serval species of fish display a huddling behavior when placed in a novel environment or are subject to stress (e.g. [Johnston, and Glasgow, 2015; Miller, 1963]). It is frequently observed in *A. lepthorhynchus* when they are introduced to a new environment. The THC-induced decrease in clustering could be the consequence of a decrease in the anxiety-like response in a novel environment.

It has been shown that chirping correlates with increased or decreased aggressiveness depending on the pattern of chirp exchanges [Hupé, 2012]. Specifically, when emitted as an echo response to a chirping conspecific, it mediates a de-escalation of aggression whereas if the chirps are not produced in response to the other fish’s chirping, it correlates with an escalation of aggression. In our experiments, the fish increased chirping as an echo-response to a stimulus containing chirps. This experiment was designed to replicate aggressive interactions and thus the increase in echo response chirping could be interpreted as a de-escalation signal. This interpretation is also supported by the decrease in aggressive lunges observed in parallel.

Our results therefore suggest that THC caused a decrease in aggressiveness and potentially a decrease in stress/anxiety and that these effects could explain the context-dependency of some of the behavioral effects observed. This hypothesis is strengthened by a rich literature on the link between the CB system and anxiety-like behaviors, including several studies on teleost species such as zebra fish. Activation of the CB system has a clear anxiolytic/anxiogenic effect in zebrafish where low or medium doses of agonist lead to decreased anxiety-like behavior but increase anxiety-like behavior at high doses [Krug, and Clark, 2015]. For example, Stewart and Kalueff [2014] tested freely swimming fish and found that, in addition to decreased mobility, high doses of THC caused a decreased in time spent in the upper portion of the tank, an anxiogenic-like response. In contrast, at low doses Barba-Escobedo and Gould [2012] described an anxiolytic-like effect of the synthetic cannabinoid WIN55,212-2 leading to more shoaling and increased exploration of lighted portions of the maze. As noted by Krug et al. [2015], the effect of CB agonists in zebrafish in various experiments depends on many variables including dose but also context.

The literature on mice show similar context-dependent effects of CB agonists. Some studies show selective decreases in aggressive behaviors while others show induction of fearfulness and inhibition of social behaviors [Cutler, and Mackintosh, 1984; Miczek, 1978; van Ree et al., 1984]. Similarly, effects of CB antagonists of mice in an elevated plus-maze, a common screen for anxiolytic drugs, showed both anxiogenic- and anxiolytic-like effects depending, on dose and context [Akinshola et al., 1999; Navarro et al., 1997]. The role of context on the influence of the CB system on anxiety and social behavior was explicitly tested in a study using CB_1_ knock-out mice. In home cages, an intruder elicited more aggressive interactions in CB_1_ deficient mice, as compared with wild-type controls, which express normal levels of CB_1_ [Haller et al., 2004]. In a novel environment, introduction of a conspecific induced fewer interactions and lower aggressiveness in knock-outs [Haller et al., 2004]. In addition, CB_1_ deletion decreased the amount of time spent in the open arms of an elevated plus maze, but only when the maze was brightly lit, indicating that CB_1_ effects on exploration are context dependent, possibly due to interactions between CB signaling and the hypothalamic-pituitary-adrenal axis [Haller et al., 2004].

Considering the widespread expression of CB receptors throughout the brain [Patel et al., 2017], it is not surprising that systemic activation/deactivation of CB receptors has complex effects that are task dependent. It is noteworthy that many of the behavioral effects described here are comparable to the effects observed in other vertebrates. Therefore, in conclusion, we argue that our study lays the basis for future studies using the advantages of the weakly electric fish as a model system to understand the dynamics of the CB system in a generalizable way.

